# *In Vivo* Phenotypic Vascular Dysfunction Extends Beyond the Aorta in a Mouse Model for Fibrillin-1 (*FBN1*) Mutation

**DOI:** 10.1101/2023.11.18.567641

**Authors:** T Curry, M.E. Barrameda, T. Currier Thomas, M. Esfandiarei

## Abstract

In individuals with Marfan Syndrome (MFS), fibrillin-1 gene (*FBN1*) mutations can lead to vascular wall weakening and dysfunction. The experimental mouse model of MFS (*FBN1^C1041G/+^*) has been advantageous in investigating MFS-associated life-threatening aortic aneurysms. Although the MFS mouse model presents an accelerated-aging phenotype in elastic organs (e.g., lung, skin), the impact of *FBN1* mutations on other central and peripheral arteries function and structure with the consideration of the impact of sex remains underexplored. In this study, we investigate if *FBN1* mutation contributes to sex-dependent alterations in central and cerebral vascular function similar to phenotypic changes associated with normal aging in healthy control mice. *In vivo* ultrasound imaging of central and cerebral vasculature was performed in 6-month-old male and female MFS and C57BL/6 mice and sex-matched 12-month-old (middle-aged) healthy control mice. Our findings confirm aortic enlargement (aneurysm) and wall stiffness in MFS mice, but with exacerbation in male diameters. Coronary artery blood flow velocity (BFV) in diastole was not different but left pulmonary artery BFV was decreased in MFS and 12-month-old control mice regardless of sex. At 6 months of age, MFS male mice show decreased posterior cerebral artery BFV as compared to age-matched control males, with no difference observed between female cohorts. Reduced mitral valve early-filling velocities were indicated in MFS mice regardless of sex. Male MFS mice also demonstrated left ventricular hypertrophy. Overall, these results underscore the significance of biological sex in vascular function and structure in MFS mice, while highlighting a trend of pre-mature vascular aging phenotype in MFS mice that is comparable to phenotypes observed in older healthy controls.

## Introduction

Marfan Syndrome (MFS) is a connective tissue disorder caused by mutations in the fibrillin-1 (*FBN1*) gene.^1,2^ The *FBN1* gene encodes for a large glycoprotein, a structural component of calcium-binding microfibrils in the extracellular matrix (ECM), providing a scaffold and structural support for elastic and non-elastic connective tissues.^3^ Therefore, *FBN1* mutations can result in a wide-range of complications in the cardiovascular, musculoskeletal, and pulmonary systems, with ascending aortic aneurysm and rupture being the most life-threatening phenotypes. MFS individuals with vascular dysfunction present weakening of blood vessel walls and progressive dilation. MFS vascular complications are attributed to compromised ECM, medial elastin fiber fragmentation, and endothelial dysfunction, all leading to vascular wall stiffening and weakening.^4–9^ Current research targets the prevention of aortic root aneurysm and rupture through lifestyle changes and pharmaceutics, but with the wide heterogeneity in *FBN1* mutations phenotype, it is necessary to consider the possibility of other associated systemic effects.

Within the ECM, FBN1 protein acts by binding the latent form of the cytokine transforming growth factor-β (TGF-β) via a latent binding protein (LTBP), forming a large sequestering domain. However, the abnormal and dysfunctional FBN1 protein fails to sequester TGF-β in its inactive form. This uncontrolled release of bioavailable TGF-β facilitates the activation of downstream signaling molecules involved in inflammatory responses, including matrix metalloproteinases (MMPs), whose role in ECM turnover involves elastin degradation.^7,10,11^ To exacerbate the dysregulation of these molecules, SMAD2/3, a downstream effector of the TGF-β signaling pathway, acts as a transcription factor and positively feedbacks to further increase TGF-β expression and bioavailability.^11^ Notably, elastin is highly expressed within the aortic wall, supporting the elasticity and distensibility of the ascending aorta. Previous studies in MFS patients and experimental mouse models have demonstrated increased expression and activity of MMP-2/-9 and significant fragmentation and disorganization of elastin fibers within the aortic media, resulting in potentially fatal outcomes due to aortic root enlargement, aneurysm, and rupture if left untreated.^5,7,10,12^ During the last two decades, the aortopathy in MFS patients and animal models has been studied in detail by many research groups. Nevertheless, our knowledge of other vascular complications in MFS is very limited. A few studies in human MFS population have reported on pulmonary artery dilation in MFS patients, as well as coronary artery dilation after aortic replacement surgery.^13–15^ Furthermore, MFS patients reportedly have an increased prevalence of cerebral aneurysms and stroke (hemorrhagic and ischemic).^16,17^ Despite the previous reports of other central and peripheral dysfunctions in MFS patients, there is no solid and comprehensive study of the structural and functional changes in other types of arteries in the well-established experimental mouse model of MFS. One translational study using the mouse model of MFS has shown increases in reactive oxygen species (ROS), MMP9, and TGF-β expression in the middle cerebral artery (MCA), all associated with an increase in wall thickness/lumen ratio measurements in MFS mice.^18^

Based on published reports, it is indicated that MFS patients and mice display accelerated and pre-mature aging phenotypes within the vasculature including increased vascular wall stiffness, decreased endothelial function, decreased smooth muscle contractility, and increased inflammatory infiltrates within the vessel wall.^6–8,10^ In addition, endothelial dysfunction associated with accelerated vascular aging can lead to decreased expression of vasodilators and anti-inflammatory molecules, resulting in increased vascular inflammation.^9,10,12^ In addition to accelerated vascular aging, MFS patients experience other age-related changes such as worsening of joint pain and stiffness, vision problems, hearing loss, skin elasticity, and lung complications.^1,2,19–21^ Evaluation of these aging-related characteristics requires further investigations, given that the mechanisms known to contribute to MFS-associated symptomology are also implicated in aging-related manifestations such as inflammation, ROS production, arterial remodeling, and elastolysis.

Incidence or prevalence of MFS does not differ by biological sex. On the other hand, MFS patients experience varied risks and complications based on sex. While female MFS patients have an increased risk of complications during pregnancy, male MFS patients have an increased risk of aortic dissection and rupture compared to non-pregnant female MFS patients.^22^ Though most preclinical MFS studies have utilized male or combined sex analysis, studies utilizing both sexes demonstrate that MFS male mice have enhanced aortic aneurysm growth, contractility, stiffness, and elastin fragmentation compared to females.^4,22–25^ Thus, evaluation of sex differences beyond the aorta is pertinent for prognosis, understanding potential mechanisms of actions, and personalized patient treatment.

In the present study, using *in vivo* ultrasound imaging, we aimed to test the hypothesis that a missense mutation in *FBN1* leads to alterations in central (aorta, pulmonary artery, coronary artery) and cerebral arteries in 6-months old MFS mice (6M-MFS) compared to sex- and age-matched controls (6M-CTRL), and that the outcomes in 6M-MFS mice are more similar to phenotypes observed in sex-matched older healthy control (12M-CTRL) mice.

## Results

### Measurements of aortic root diameters and wall stiffness in MFS mice

Ultrasound imaging was utilized to assess aortic root diameters between MFS and CTRL (**Fig. 1A & 1B**) as a function of sex (**Fig. 1C-D)** and age (**Fig. 1F-K**). Increased aortic root diameters are indicative of aortic root enlargement. The aortic annulus diameter was increased by 36% in 6M-MFS male mice compared to 6M-CTRL male mice, and by 18% compared to age-matched MFS female mice (**Fig. 1C**). At the sinus of Valsalva, both male (29%) and female (21%) 6M-MFS mice display an increase compared to sex- and age-matched CTRL mice without sex differences (**Fig. 1D**). At the sinotubular junction, 6M-MFS male mice demonstrated an increase of 32% compared to 6M-CTRL male and of 34% compared to 6M-MFS female mice (**Fig. 1E**).

**Figure 1.**
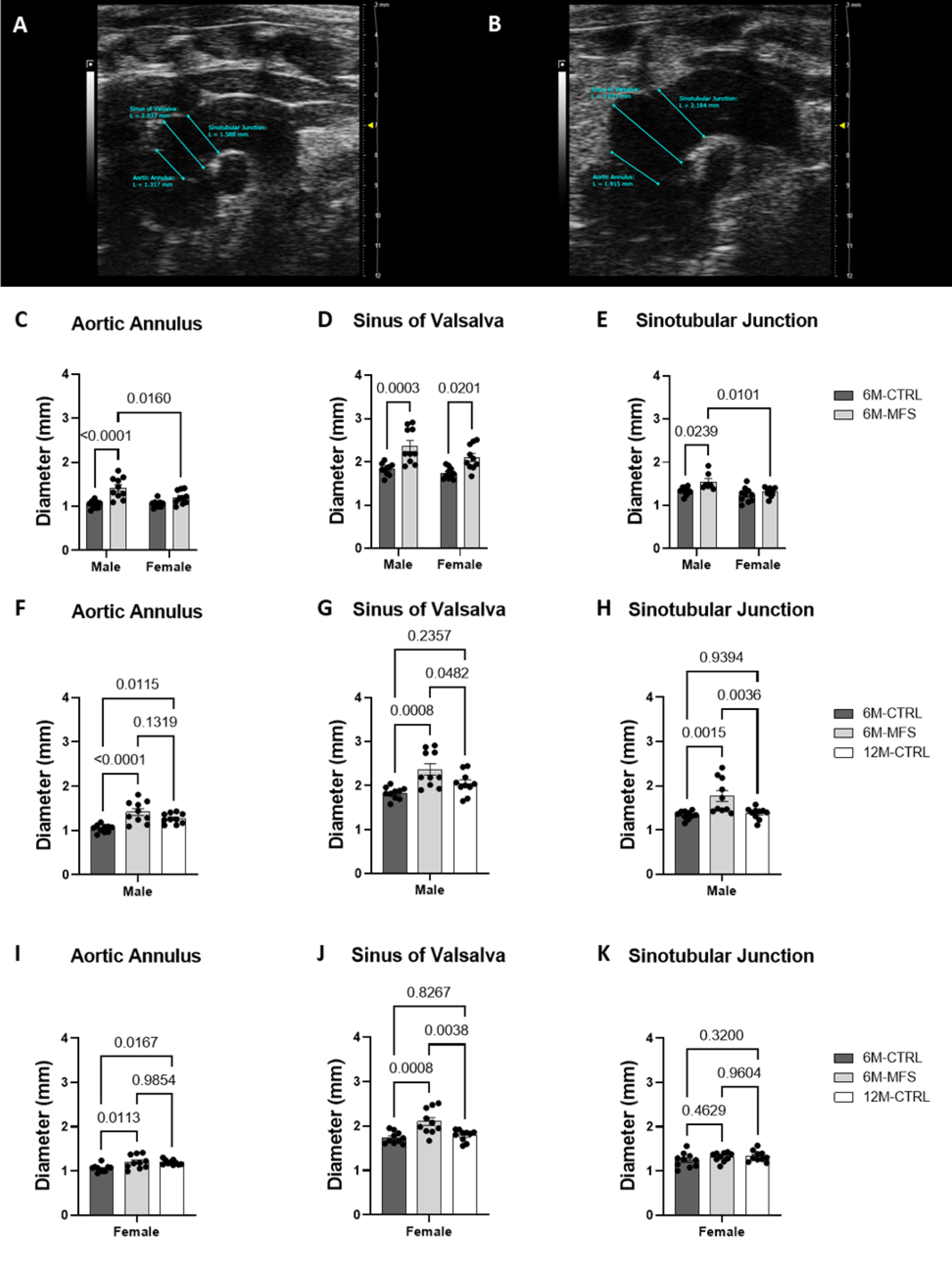
Measurements of aortic root diameters in MFS mice. **A)** Representative B-mode image of the aortic arch of a 6M-CTRL male evaluated for diameters of the aortic annulus, sinus of Valsalva, and sinotubular junction. **B)** Representative B-mode image of the aortic arch of a 6M-MFS male with aortic diameters analyzed. **C)** 6M-MFS male aortic annulus diameters were increased compared to 6M-CTRL male mice as well as 6M-MFS female mice. **D)** 6M-MFS male and female mice demonstrate increased sinus of Valsalva diameters compared to sex- and age-matched CTRL mice. **E)** 6M-MFS male sinotubular junction diameters were increased compared to 6M-CTRL male mice as well as 6M-MFS female mice. **F)** In the Aortic Annulus, 6M-MFS and 12M-CTRL male mice have increased diameter compared to 6M-CTRL male mice. 6M-MFS male mice have increased **G)** Sinus of Valsalva and **H)** Sinotubular Junction diameters compared to 6M- and 12M-CTRL males. **I)** 6M-MFS females and 12M-CTRL females have increased Aortic Annulus diameters compared to 6M-CTRL female mice. **J)** 6M-MFS females have increased Sinus of Valsalva diameters compared to 6M- and 12M-CTRL female mice. **K)** No differences were seen in the Sinotubular Junction diameter of our female cohorts.

We also compared aortic diameters in all three regions of the aortic root between 6M-CTRL, 6M-MFS, and 12M-CTRL of each sex to determine whether the MFS phenotype was more similar to older mice, rather than age-matched controls. In males, 6M-MFS and 12M-CTRL mice had similar increases in aortic annulus diameters compared to 6M-CTRL male mice (**Fig. 1F**) but differed at the sinus of Valsalva and sinotubular junction diameters (**Fig. 1G & 1H**). In female mice, 6M-MFS and 12M-CTRL mice had similar increases in aortic annulus diameters compared to 6M-CTRL female mice (**Fig. 1I**) but differed at the sinus of Valsalva (**Fig. 1J**) and showed no differences at the sinotubular junction between the female mice (**Fig. 1K**).

Aortic pulse wave velocity (PWV) is a well-established measure of aortic wall stiffness. Increased PWVs are indicative of elevated wall stiffness and vascular rigidity, which is a hallmark of vascular pathology in MFS patients and experimental mouse models, making the ascending aorta more vulnerable to aortic root dissection and rupture.^6–8^ We utilized ultrasound imaging (color Doppler) to measure PWV in sex and age-matched CTRL and MFS mice (**Fig. 2A-C**). Our data supports previous reports showing an increase in aortic PWV in 6M-MFS mice compared to sex- and age-matched CTRL mice (**Fig. 2D**). A significant difference in PWV values was observed in 6M-old male MFS as compared to sex-matched 12M-old healthy CTRL mice (**Fig. 2E**), indicating this outcome measure is unique to the genotype. On the other hand, PWV values in 6M-old female MFS mice are comparable to those measured in sex-matched 12M-old mice (**Fig. 2F**), demonstrating that only female MFS mice show aging-like aortic phenotypes.

**Figure 2.**
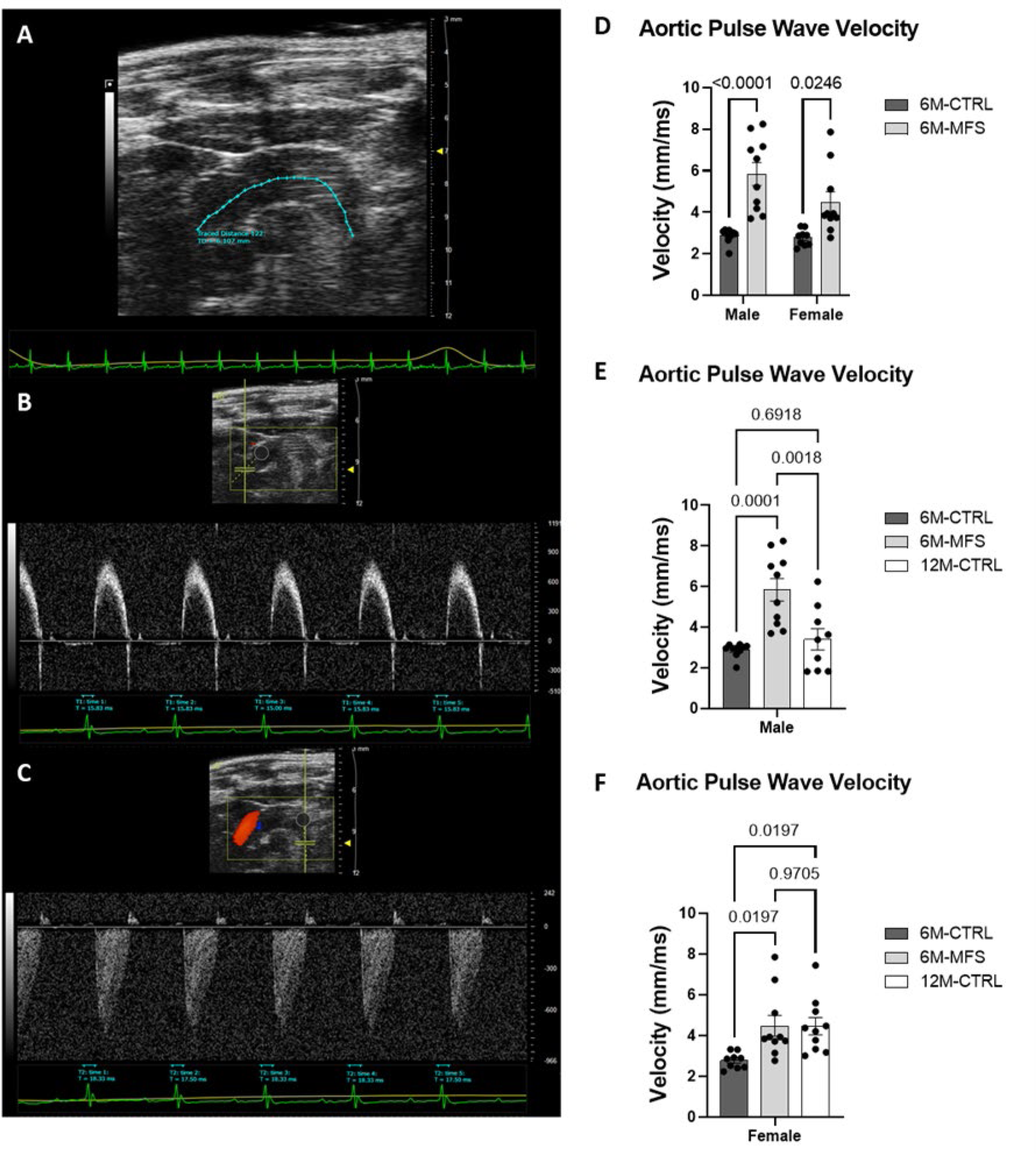
Measurements of aortic root wall stiffness in MFS mice. **A)** Representative image of a 6M-CTRL female traced aortic arch in B-mode. **B)** PW Doppler Mode waveform of the ascending aortic arch (Time 1, T1) with representative times analyzed. **C)** PW Doppler Mode waveform of the descending aortic arch (Time 2, T2) with representative times analyzed. D) 6M-MFS male and female mice have increased aortic stiffness, compared to sex-& age-matched CTRL mice. **E)** 6M-MFS male mice have increased aortic stiffness, measured as aortic pulse wave velocity, compared to 6M- and 12M-CTRL male mice. **F)** 6M-MFS female and 12M-CTRL female mice have increase aortic pulse wave velocity compared to 6M-CTRL female mice.

### *Measurements of* coronary artery peak blood flow velocity in MFS mice

To investigate the impact of MFS on coronary artery function, we used *in vivo* ultrasound imaging to measure the peak blood flow velocity of the coronary artery between CTRL and MFS mice as a function of sex and age (**Fig. 3A**) No difference was seen in coronary artery peak blood flow velocity during diastole in male or female MFS mice as compared to sex- and age-matched CTRL mice (**Fig. 3B**). Furthermore, sex differences were not demonstrated in coronary blood flow velocity (**Fig. 3B**). Peak blood flow velocity was also similar between 6M-CTRL, 6M-MFS, and 12-CTRL in males (**Fig 3C**) and females (**Fig. 3D**).

**Figure 3.**
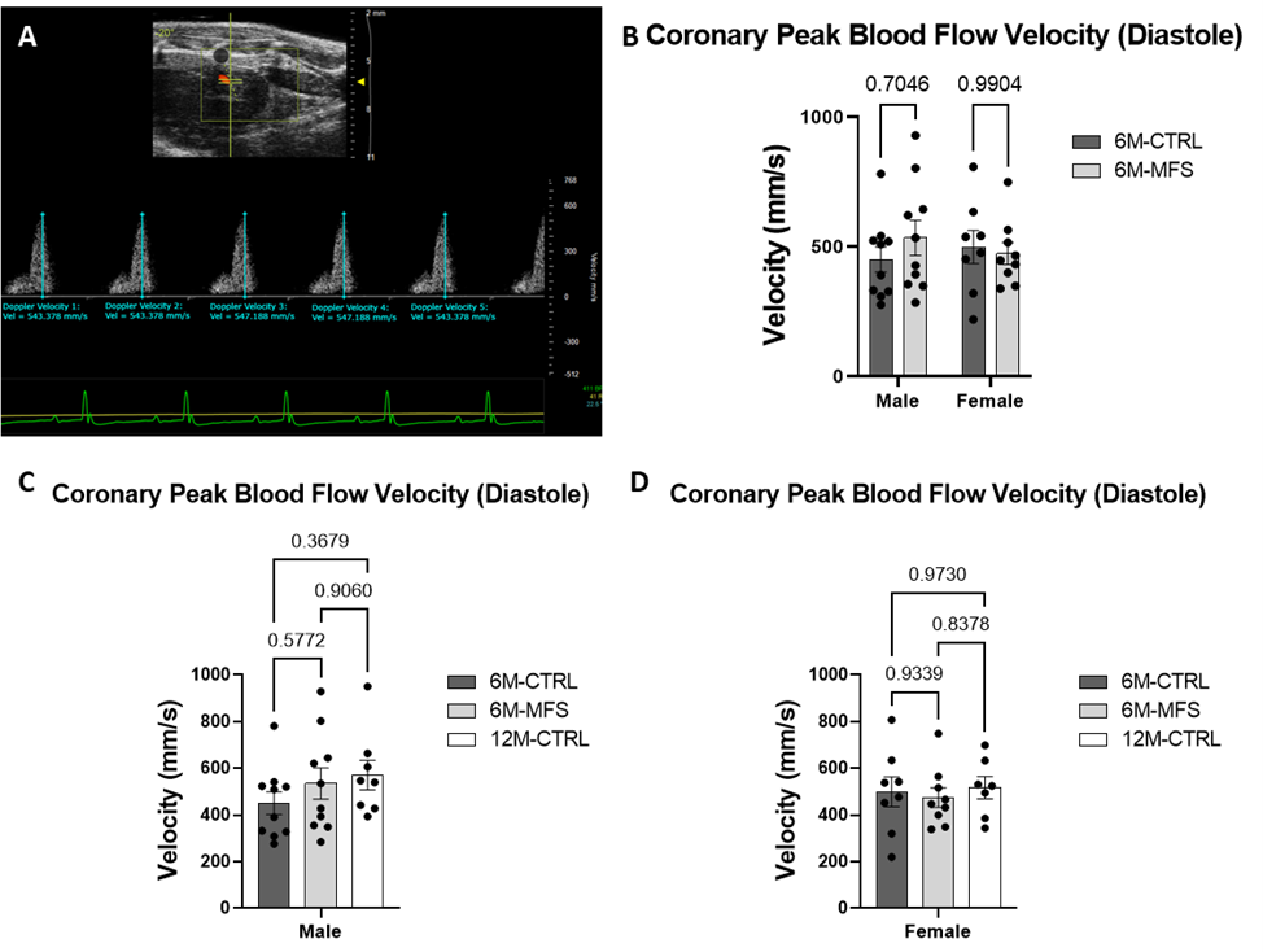
Measurements of coronary artery peak blood flow velocity during diastole in MFS mice. **A)** Representative image of coronary artery outflow visualized in the parasternal long axis view with pulse wave doppler mode waveform analyzed with five consecutive velocity measurements in a 6M-CTRL female mouse. **B)** Genotype and sex did not induce differences in coronary peak blood flow velocity during diastole at 6M. No differences were seen in coronary peak blood flow velocity during diastole as a function of age in **C)** male cohorts or **D)** female cohorts.

### Assessments of left pulmonary artery peak blood flow velocity in MFS mice

To investigate pulmonary artery function, peak blood flow velocity was evaluated *in vivo* between CTRL and MFS mice as a function of sex and age (**Fig. 4A**). 6M-MFS male and female mice demonstrated significantly lowered peak blood flow velocity in the left pulmonary artery as compared to sex- and age-matched CTRL groups (**Fig. 4B**). Sex differences were not demonstrated in pulmonary blood flow velocity measurements (**Fig. 4B**). Blood flow velocity values in 12M-CTRL male and female mice were also decreased compared to sex-matched 6M-CTRL mice, without differences as compared to 6M-MFS mice (**Fig 4C & 4D**), indicating a decrease in pulmonary blood flow velocity due to genotype and aging.

**Figure 4.**
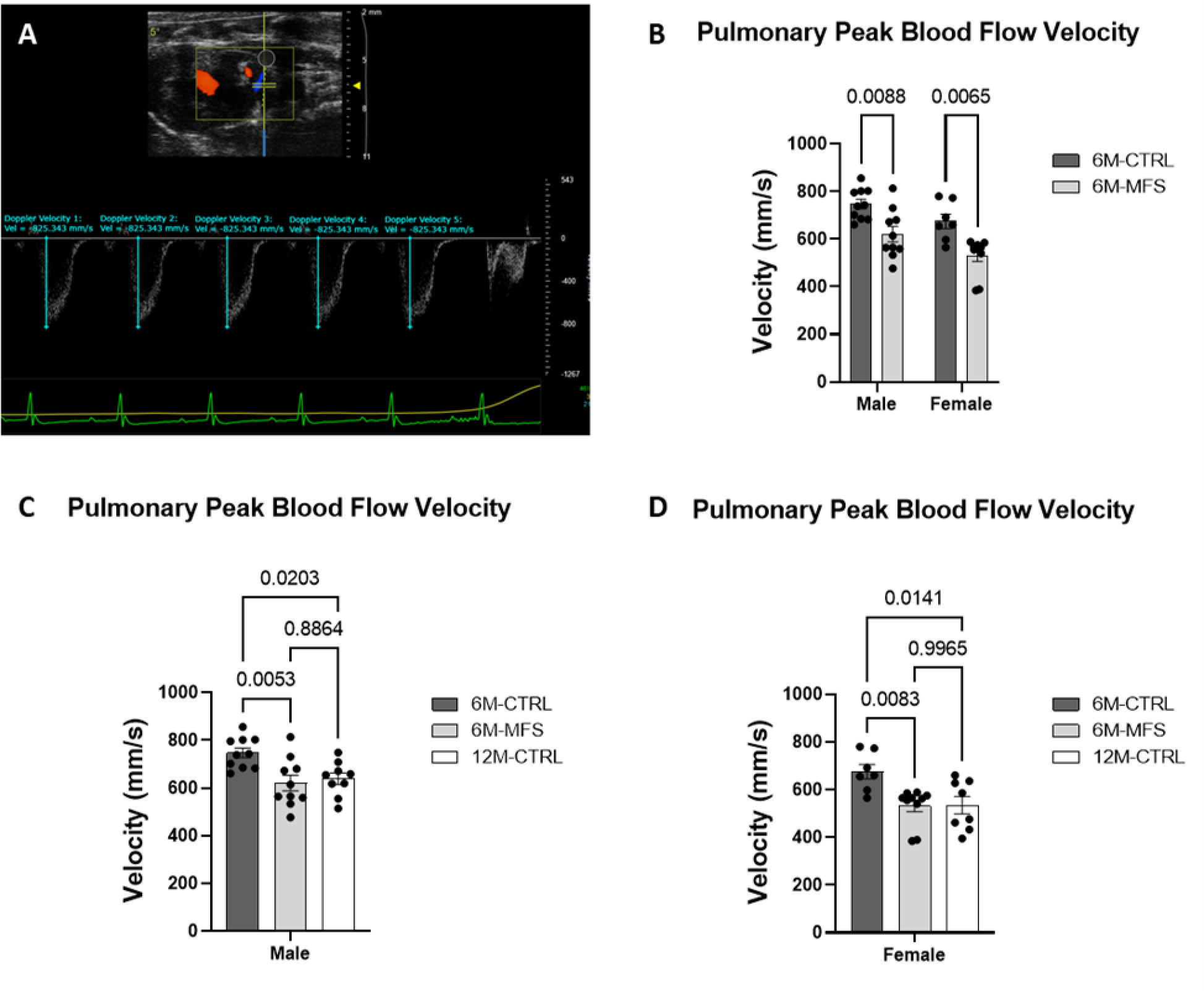
Assessments of left pulmonary artery peak blood flow velocity in MFS mice. **A)** Representative image of left pulmonary artery outflow visualized in the parasternal long axis view with pulse wave doppler mode waveform analyzed with five consecutive velocity measurements in a 6M-CTRL male mouse. **B)** 6M-MFS male and females have decreased left pulmonary artery peak blood flow velocity compared to sex- and age-matched 6M-CTRL mice. 6M-MFS and 12M-CTRL mice have decreased left pulmonary peak blood flow velocity compared to 6M-CTRL in **C)** male and **D)** female cohorts.

### Measurements of posterior cerebral artery peak blood flow velocity in MFS mice

Our understanding of the alterations in structure and function of cerebral arteries in the MFS mouse model remains incomplete. In this study, we utilized ultrasound imaging Doppler analysis to assess posterior cerebral artery (PCA) peak blood flow velocity between CTRL and MFS mice as a function of sex and age (**Fig. 5A & 5B**). 6M-MFS male mice have decreased PCA peak blood flow velocity compared to sex- and age-matched CTRL mice (**Fig. 5C**) with no difference observed between age-matched female MFS and CTRL (**Fig. 5C**). Pertinently, 6M-CTRL female mice also demonstrated lower PCA peak blood flow velocity compared to age-matched male CTRL mice, highlighting a fundamental sex difference in this outcome measure (**Fig. 5C**). As shown in **figure 5D**, outcomes of 6M-MFS male and 12M-CTRL male mice are similar, displaying a significantly decreased PCA blood flow velocity compared to 6M-CTRL male mice. In females, age did not influence PCA peak blood flow velocity (**Fig. 5E**).

**Figure 5.**
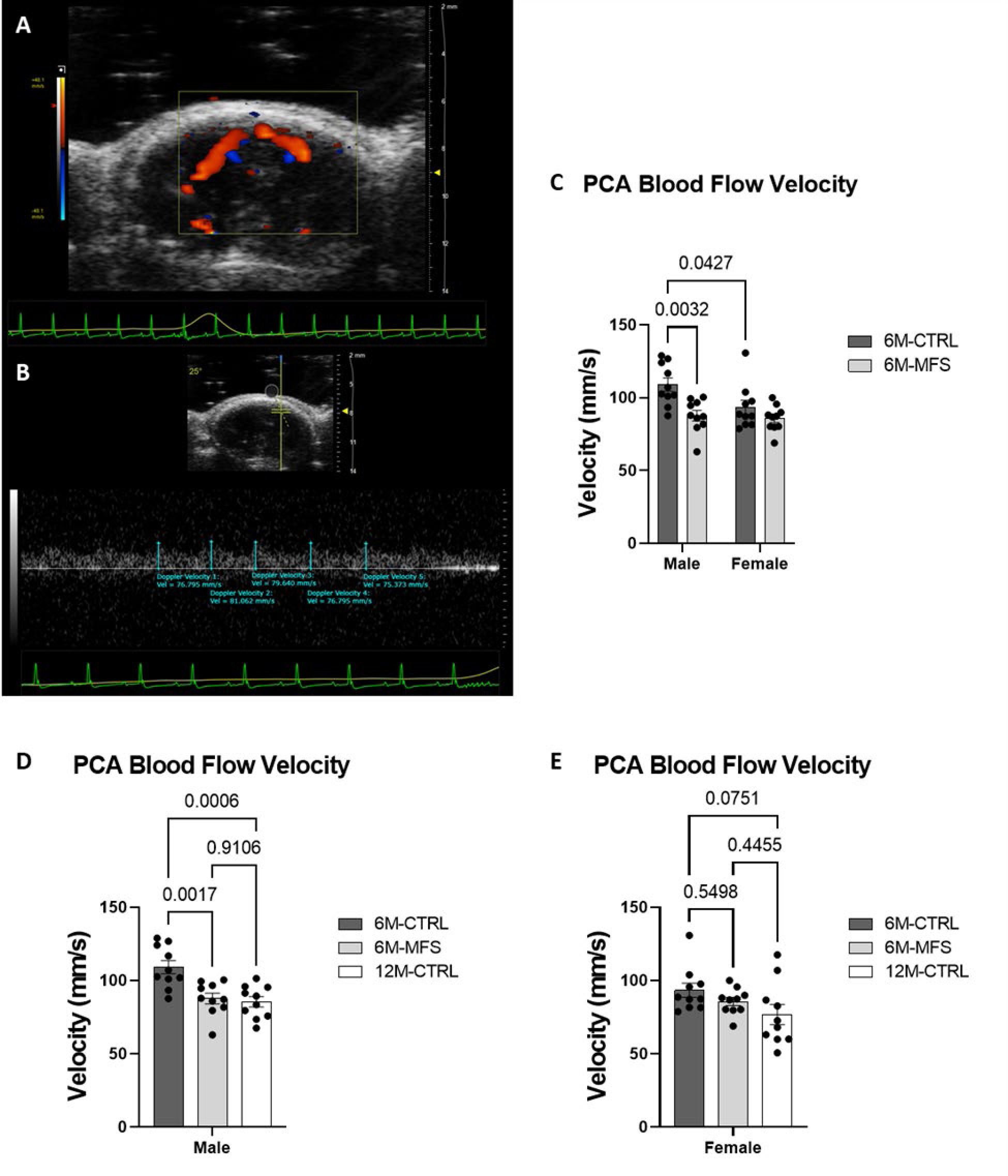
Measurements of posterior cerebral artery peak blood flow velocity in MFS mice. **A)** Representative image of color doppler mode of the PCA in a 6M-CTRL female. **B)** Representative image of Pulse Wave Doppler Mode waveform of the PCA analyzed with five consecutive velocity measurements. **C)** 6M-MFS male mice and 6M-CTRL female mice demonstrated decreased PCA blood flow velocity compared to 6M-CTRL male mice. **D)** 6M-MFS and 12M-CTRL male mice have decreased PCA blood flow velocity compared to 6M-CTRL mice. **E)** Female mice showed no differences amongst cohorts.

### Comprehensive evaluation of cardiac function and structure in MFS mice

*In vivo* ultrasound imaging was used to evaluate cardiac parameters including cardiac output, stroke volume, heart rate, ejection fraction, fractional shortening, and left ventricular mass (**Fig. 6A & 6B & 6C**). Our data indicates that by 6-months of age MFS in mice is not associated with any significant change in the cardiac output, stroke volume, heart rate, ejection fraction, or fractional shortening compared to age-matched CTRL mice in either male or female groups (**Tables 1 & 2**). Contrarily, left ventricular mass was significantly increased in 6M-MFS male mice compared to sex- and age-matched CTRL mice with no difference observed when compared to 12M-CTRL male mice (**Table 1**). However, no differences in left ventricular mass were observed in female 6M-MFS mice when compared to sex- and age-matched CTRL mice (**Table 2**).

**Figure 6.**
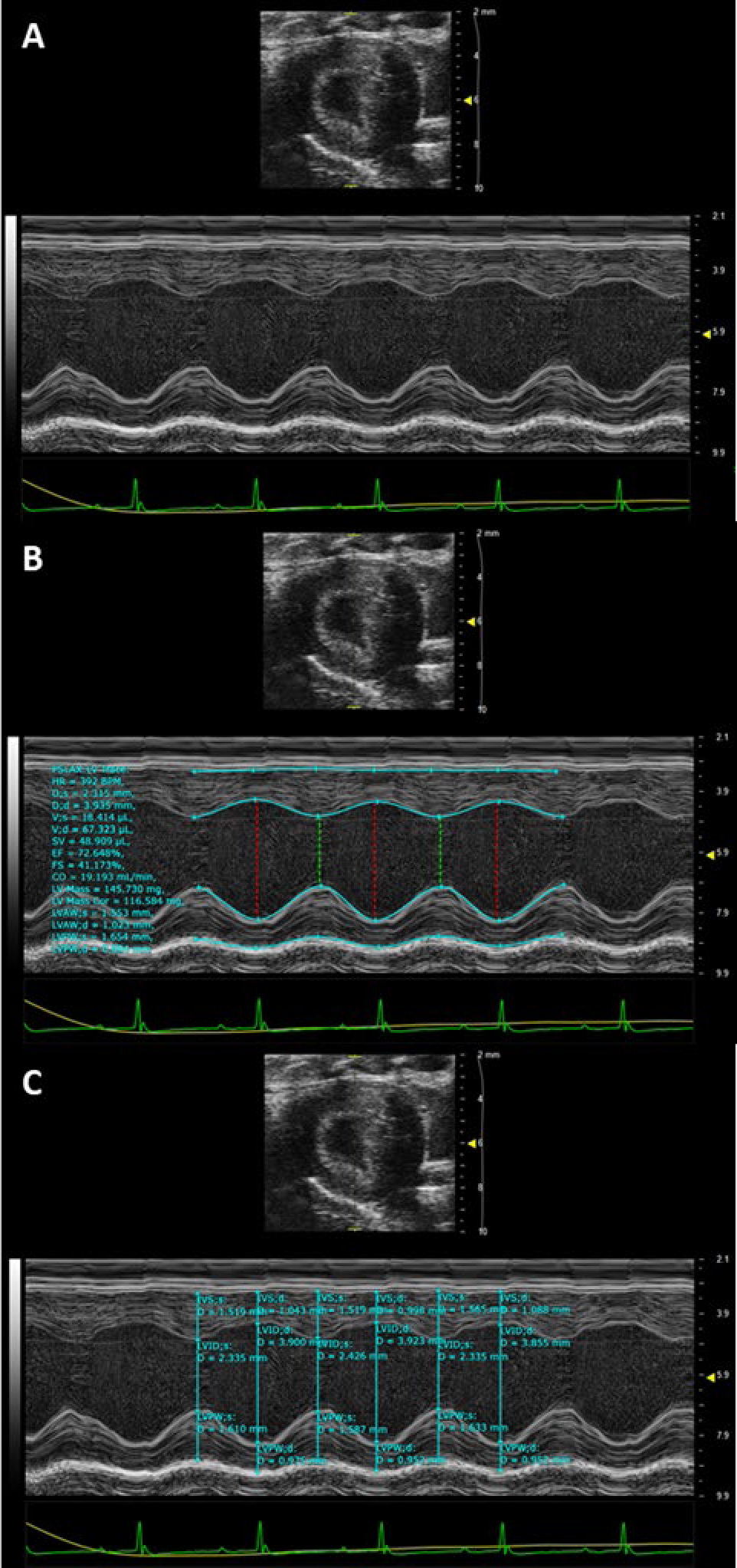
Representative M-mode images of parasternal long axis view to visualize the left ventricle in a 6M-CTRL male mouse. **A)** The left ventricle tracing of the anterior and posterior walls is visualized. **B)** Parasternal long axis left ventricle trace (PSLAX LV trace) analysis allows for tracing of the ventricle over three consecutive contractions to measure cardiac parameters and **C)** intraventricular wall septum in systole and diastole.

**Table 1.** Comprehensive evaluation of cardiac function and structure in MFS male mice. Values are mean ± standard error of the mean (SEM). One-way ANOVA p-value displayed. Significant differences for multiple compares demonstrated as *Compared to 6M-CTRL, #Compared to 6M-MFS, and †Compared to 12M-CTRL.

| Male |  |  |  |  |
| --- | --- | --- | --- | --- |
|  | 6M-CTRL | 6M-MFS | 12M-CTRL | One-way ANOVA p-value |
| Weight (g) | 38.20±1.428† | 41.78±1.956 | 47.60±2.500* | 0.009 |
| Heart Rate (beats/min) | 435.8±14.95 | 437.1±12.22 | 449.7±11.94 | 0.742 |
| Stroke Volume (μl) | 45.37±1.201 | 46.48±4.837 | 51.21±2.270 | 0.451 |
| Cardiac Output (mL/min) | 19.71±0.681 | 20.06±1.953 | 23.12±1.449 | 0.241 |
| Ejection Fraction (%) | 73.01±2.415 | 74.09±3.716 | 77.25±2.199 | 0.603 |
| Fractional Shortening (%) | 41.85±2.002 | 43.52±3.280 | 45.79±2.227 | 0.591 |
| Left Ventricular Mass (mg) | 119.6±3.371# | 174.2±16.15* | 146.4±12.70 | 0.005 |
| Mean ± SEM *Compared to 6M-CTRL, #Compared to 6M-MFS, †Compared to 12M-CTRL | | | | |

**Table 2.** Comprehensive evaluation of cardiac function and structure in MFS female mice. Values are mean ± standard error of the mean (SEM). One-way ANOVA p-value displayed. Significant differences for multiple compares demonstrated as *Compared to 6M-CTRL, #Compared to 6M-MFS, and †Compared to 12M-CTRL.

| Female |  |  |  |  |
| --- | --- | --- | --- | --- |
|  | 6M-CTRL | 6M-MFS | 12M-CTRL | One-way ANOVA p-value |
| Weight (g) | 31.00±1.976 | 30.00±2.206† | 37.50±1.918# | 0.039 |
| Heart Rate (beats/min) | 457.3±10.93 | 450.1±8.787 | 446.5±9.895 | 0.735 |
| Stroke Volume (μl) | 42.23±3.175 | 35.51±3.865 | 31.08±3.791 | 0.103 |
| Cardiac Output (mL/min) | 19.21±1.313 | 15.99±1.800 | 13.71±1.504 | 0.054 |
| Ejection Fraction (%) | 71.74±1.845 | 79.91±2.298 | 77.58±3.383 | 0.263 |
| Fractional Shortening (%) | 40.48±1.562 | 47.83±2.104 | 45.82±2.869 | 0.065 |
| Left Ventricular Mass (mg) | 115.3±6.265 | 131.2±10.30 | 134.1±8.998 | 0.253 |
| Mean ± SEM *Compared to 6M-CTRL, #Compared to 6M-MFS, †Compared to 12M-CTRL | | | | |

Using ultrasound imaging, we also assessed the mitral valve early filling (MV E) wave velocity, mitral valve atrial (MV A) filling wave velocity, and the E/A ratio of these velocities (**Fig. 7A**). Our findings indicate that at 6 months of age, MFS males and females demonstrate decreased MV E velocities compared to sex- and age-matched CTRL mice without sex differences seen (**Fig. 7B**). No differences in MV A velocity or E/A ratio values were observed across the experimental groups (**Fig. 7C & 7D**). Age did not contribute to differences amongst male MV E, MV A, or E/A ratio measurements (**Fig. 7E-7J**). The values for MV E were significantly decreased in 6M-MFS male and female mice as compared to sex- and age-matched CTRL mice (**Fig. 7E & 7H**). The significant drop in MV E values in 6M-MFS females were comparable with those observed in 12M-CTRL female mice, highlighting an accelerated aging phenotype in young female MFS mice (**Fig. 7H**). However, we did not make the same observation between 6M-MFS and 12M-CTRL male mice (**Fig. 7E**). No differences were demonstrated amongst the female cohorts in the MV A or E/A ratio measurements (**Fig. 7I & 7J**).

**Figure 7.**
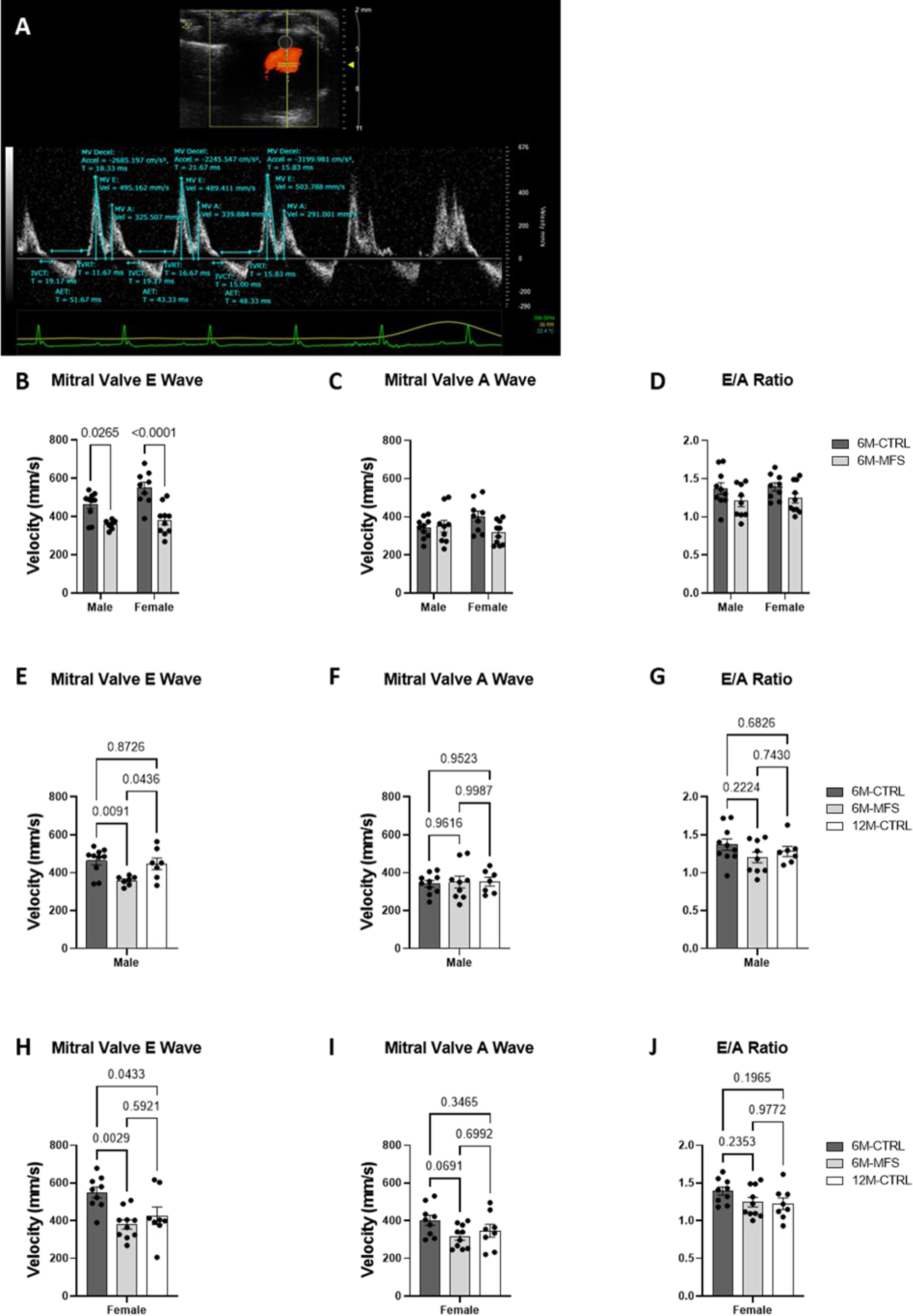
Comprehensive evaluation of mitral valve function in MFS mice. **A)** Representative Image of Pulse wave Doppler Mode of the mitral valve inflow while in the apical four chamber view with measurements for peak early filling (MV E), peak atrial filling (MV A), and E/A ratio in a 6M-CTRL male mouse. **B)** Male and female 6M-MFS mice have decreased MV E filling velocity compared to sex-and age-matched CTRL mice but without differences in **C)** MV A filling velocity or **D)** E/A ratio. **E)** MV E filling velocity is decreased in 6M-MFS males compared to both 6M- and 12M-CTRL male mice. No differences were demonstrated in **F)** MV A filling velocity or **G)** E/A ratio in the male cohorts. **E)** MV E filling velocity was decreased in 6M-MFS and 12M-CTRL female mice compared to 6M-CTRL female mice. **F)** MV A filling velocity and **G)** the E/A ratio is unchanged in female mice.

## Discussion

Prior studies utilizing MFS mouse models have focused on the life-threatening manifestation of aortic aneurysm; thus, few studies have evaluated *in vivo* alterations in other arteries, particularly with the consideration for sex differences in experimental subjects. The accelerated aging phenotype is of interest as it offers unique insight into fundamental processes underlying vascular aging with potential relevance for TGF-β’s signaling role in vascular aging or for potential therapeutic targets. Mostly, we exhibited that male MFS mice had more robust genotypic changes than females, and these genotypic changes tended to be more comparable to middle-aged (12M-old) healthy sex-matched CTRL mice than those recorded in 6M-old sex-matched CTRL mice.

This study is the first in-depth examination of structural/functional changes in central arteries (including aorta, pulmonary, coronary) and cerebral arteries in the well-established mouse model of MFS, with the consideration of sex as a distinct biological variable. Though there is no sex-related difference in the incidence of MFS, male patients experience an increased risk for aortic dissection and rupture.^22^ In MFS mice, it has also been demonstrated that male subjects have enhanced aortic root aneurysm growth compared to females.^22–25^ Aortic contractility and stiffness also demonstrate sex differences in MFS mice, where MFS males display higher smooth muscle contractility and aortic wall elastin fragmentation compared to MFS females.^4^ Further evaluation of contractility and relaxation utilizing myography to asses sex differences would be paramount in understanding the mechanisms behind these differences.^26^ Sex-related differences in the aorta have been associated with increased mitogen-activated protein kinase ERK1/2 phosphorylation, and enhanced activity of endothelial nitric oxide synthase (eNOS) in females, underscoring sexual dimorphism and estrogen-associated endothelial protection.^4,22,23^ Investigating sex-based differences is critical for a deeper understanding of potential variables that contribute to designing care plans for MFS patients through more individually tailored strategies. Thus, our analysis evaluated sex as a biological variable throughout the described measures.

The current approaches in MFS management and treatment have extended the life expectancy of affected individuals from approximately 40 to around 75 years of age.^27,28^ As a result, this population could be more susceptible to other vascular complications commonly associated with aging. Consequently, there is a need for further research to explore other symptoms and quality of life (QoL) issues in patients with MFS and related connective tissue disorders. Primary complaints contributing to lowered QoL in this patient population include pain, mobility issues, fatigue, and anxiety/depression.^19,28–30^ Understanding how MFS affects the vascular system over time will help clinical scientists and clinicians design more effective preventive and therapeutic approaches, particularly for aging MFS patients at higher risk for vascular complications. Few studies have addressed this vascular aging-associated phenotype in MFS mice. One study in MFS mice demonstrated a pre-mature aging phenotype in skin morphology.^21^ The observed mitral valve prolapse in MFS patients and experimental mouse models has been correlated with elastin degradation similar to aging-related elastolysis, where elastin degradation has been well-established to contribute to normal skin and vascular aging.^31^ Another study evaluated *FBN1* mutation associated elastin fragmentation over time in the aorta of MFS mice, where MFS mice indicated pathological aging.^32^ In this study, we aimed to investigate the pre-mature and accelerated aging phenomenon in the vasculature in MFS mice by comparing *in vivo* vascular measurements in young (6M-old) MFS to sex- and age-matched CTRL and sex-matched 12M-old CTRL mice.

In agreement with previous published work by our group and others, our results from this study also support aortic root dilation and loss of wall elasticity in MFS mice, with more prominent changes in MFS male mice. Sex differences in aortic diameters have been reported in the general population, where women tend to have smaller aortic diameters than men.^22^ To the best of our knowledge these are the first comparisons evaluating sex differences in preclinical assessments of *in vivo* aortic diameters in this mouse model. Our data in mice support that both MFS male and female mice demonstrate increased dilation and risk of aortic stiffening compared to sex- and age-matched CTRL mice, but the detrimental changes are exacerbated in MFS male mice. MFS males demonstrated increased aortic diameter throughout the ascending aorta and increased aortic PWV, a supported measure of arterial wall stiffening. Female MFS mice only demonstrated increased aortic diameters in the aortic annulus and sinus of Valsalva, which was associated with increased aortic wall stiffness. MFS males demonstrated increased diameters in the aortic annulus and sinotubular junction compared to MFS female mice. These observations suggest that future studies targeting aortic stiffness and contractility should evaluate sex as a biological variable in this mouse model. MFS mice did not show similarity to middle-aged CTRL mice in the aortic measures. This could be attributed to the aorta’s inherent elasticity, suggesting that MFS-associated elastin fragmentation and disorganization may exceed those associated with normal aging, inducing further vulnerability.

Though no differences were demonstrated in coronary blood flow velocity in diastole amongst the experimental groups, MFS patients are reported to develop with coronary artery dilation that risks aneurysm and rupture, particularly with complications after aortic replacement surgery.^15^ It is recommended that coronary artery size be regularly monitored in MFS patients, especially those with severe aortic phenotypes or after aortic replacement surgery.^15^ Thus, future studies evaluating coronary blood flow velocity during systole, as well as to assess preclinical *ex vivo* analysis of the coronary arteries evaluating stiffness and elasticity in MFS mice are warranted to further assess this risk for coronary dilation. Furthermore, this change in function may not be detectable in coronary blood flow velocity during diastole by 6-months of age in MFS mice but may be prevalent at later time points.

Pulmonary artery dilation has been documented in MFS patients and experimental mouse models and been correlated with more severe disease of the ascending aorta in MFS, warranting further investigation.^13,14,33^ A preclinical assessment of pulmonary artery dilation in MFS mice showed that by 12 weeks of age MFS mice had increased pulmonary artery diameters compared to aged-matched control mice.^13^ Aging alone has also been associated with pulmonary artery dilation, stiffening, and hypertension in clinical and preclinical assessments.^34–36^ Pulmonary dilatation would increase the blood flow throughout the pulmonary system due to a decrease in resistance. Inversely, blood flow velocity (mm/s) decreases due to the relationship between blood flow (mL/s) and cross-sectional area of the blood vessel, where velocity is the distance of the blood flow over time (Flow = Velocity * Cross-sectional Area).^37^ Thus, as the diameter or cross-sectional area of the vessel increases, the velocity decreases. This relationship is in alignment with the dilatation expected in MFS and aging. Pulmonary artery dilatation has been associated with both MFS and pulmonary hypertension, but in MFS patients the pulmonary artery enlargement is not associated with hypertension and does not commonly produce pathological symptoms related to pulmonary hypertension.^14,38^ Furthermore, this inverse relationship between diameter and velocity has been validated in humans.^39^ Blood flow velocity can also be affected by viscosity and blood pressure, which were not evaluated in this study but have not been shown to be different in MFS. Our data confirms a decrease in peak blood flow velocity in the left pulmonary artery of 6M-MFS mice that is comparable to values reported in 12M-CTRL mice. It is noteworthy that our data also shows age-associated decline, where 12M-CTRL male and female mice show a significantly lower left pulmonary artery peak blood flow velocity as compared to younger sex-matched CTRL mice, which is in agreement with previous reports in older human population.^34^

MFS patients have demonstrated an increased risk for cerebrovascular disease and hospitalization, with increased odds of ischemic stroke, carotid artery dissection, and cerebral aneurysm and intracranial hemorrhage.^17^ Overall, our results confirm these findings and indicate that MFS male mice demonstrate an accelerated cerebrovascular aging phenotype. Blood flow to the brain is supplied by the aorta, to the two internal carotid arteries, and then further into the Circle of Willis. The posterior cerebral artery (PCA) is another one of the major arteries that supply blood to the brain, it is reasonable to consider it and the associated sex differences as we extend studies into potential vulnerability of the cerebrovasculature in MFS. Here, we have shown that PCA peak blood flow velocity is significantly decreased in 6M-MFS male mice when compared to sex- and age-matched CTRL mice.

A sex difference was detected between PCA blood flow velocity in 6M-CTRL female mice compared to age-matched CTRL male mice, with females demonstrating lowered velocity than males. However, such differences were not observed between female 6M-MFS and female 6M-CTRL mice, indicating a sex-dependent effect of PCA blood flow velocity, which requires further investigation. In a prior study, the middle cerebral artery (MCA) was determined to have increased lumen/wall ratio in 6M-MFS mice compared to age-matched controls in a combined sex cohort, suggesting dilation.^18^ This finding correlates with our result of decreased blood flow velocity as described above. Furthermore, it was reported that apolipoprotein E (*ApoE*)-deficient mice with a *FBN1* mutation (*ApoE*^−/−^*FBN1*^C1041G/+^) presented with accelerated blood-brain barrier degradation, increased inflammatory cytokines (IL-1β, TNF-α), MMPs −2/−9, and TGF-β in the MCA and choroid plexus.^40,41^ Increased MMP-2/-9 expression and activity could result in increased neuroinflammation and can, directly and indirectly, alter neuronal function and neuroplasticity.^42^ There is an increased prevalence of headaches and migraines in MFS patients, suggesting cerebrovascular dysfunction.^43,44^ MFS patient case studies also report neurodevelopmental disorders, including attention deficit hyperactivity disorder (ADHD) and learning disabilities.^45–48^

In addition, connective tissue disorders such as Ehlers-Danlos syndrome have been implicated in increased vulnerability to neurological insult such as traumatic brain injury and higher rates of complication and mortality after neuroendovascular procedures.^49–51^ Decreased cerebral blood flow velocity correlated with the function of age, increases risks for severity after neurological insult, neuropathology, and cognitive impairment.^52–56^ Due to recent findings and the overlapping characteristics displayed with aging, it is plausible that the cerebral vasculature and brain could be affected in MFS patients. Together with the increased risk for cerebral aneurysms and stroke (hemorrhagic and ischemic) in MFS patients, further research is needed to understand the magnitude of risk, the potential discovery of novel targets for prevention and treatment, and the impact of current treatments on neurological concerns.^16,17,57^

We observed increased left ventricular hypertrophy in 6M-MFS male mice compared to sex- and age-matched CTRL mice but did not demonstrate this difference in females. This finding of left ventricular remodeling where the septum and posterior walls were thicker in MFS mice has been reported before but has not been evaluated for sex as a variable.^58^ The result of this study further underscores the importance of sex-based differences in vascular aging. This presented change in left ventricular mass did not correlate with altered ejection fraction by 6 months of age but could result in altered cardiac function as the animal ages. In addition, other cardiac parameters such as cardiac output, stroke volume, fractional shortening, and heart rate were not different in MFS male or female mice compared to age-matched, sex-matched, or middle-aged (12M-old) control mice.

Male and female MFS mice displayed decreased mitral valve early-filling velocities, without statistically significant differences in the E/A ratio. MFS females showed similarity to middle-aged CTRL females, suggesting an effect of genotype and aging. Though statistical difference was not demonstrated in the E/A ratio, it is clinically relevant to recognize that decreased MV E filling velocities did reduce the overall E/A ratio. In the MFS population, Mitral valve prolapse (MVP) is prevalent and shown to be more common in females compared to males, where studies have indicated that women with MVP often experience more severe symptoms and are at higher risk for complications such as mitral regurgitation and endocarditis.^59–61^

Due to evident phenotypic differences in the mice, the ultrasound technician that performed image acquisition was not blinded to genotype, and sex. To mitigate this concern, the images gathered from the imaging were de-identified, coded, and analyzed by two blinded independent investigators. Data reported are the averages of the two investigators. In the case of difficult imaging, not all measurements could be taken for each animal. All data display the individual data points to help discern this in data interpretation.

In conclusion, this study highlighted alterations in central, and cerebral vascular function that was sex dependent. Furthermore, the observed similarities in vascular functions between young MFS mice and middle-aged CTRL mice suggest the possibility of pre-mature and accelerated aging of the vasculature in MFS mice.

## Materials and methods

### Experimental Animals

Animal care was conducted according to the National Research Council Guide for the Care and Use of Laboratory Animals and the Guidelines for Animal Experiments and the Institutional Animal Care and Use Committee [IACUC protocols AZ-3006, AZ-2989, & AZ-2936]. All experiments were in compliance with ARRIVE guidelines. Mice were group-housed (up to 5 mice per cage) in a 12/12h light-dark cycle with food and water available *ad libitum*. Breeding pairs were obtained from Jackson Laboratory and backcrossed for 8 generations.

A breeding colony of MFS mice was established and maintained. These mice were heterozygous for an *FBN1* allele encoding a cysteine substitution in the epidermal growth factor-like domain in *FBN1* to glycine (*FBN1^C1041G/+^*), giving the mice the most common type of mutation seen in MFS patients presented with vascular dysfunction and aortic root aneurysm. This study utilized adult *FBN1^C1041G/+^* (MFS) mice at 6 months of age, as well as *C57BL/6* (CTRL) mice at 6 and 12 months of age. MFS mice present established and evident vascular defects (e.g., aortic wall elastin fragmentation, aortic root aneurysm, and increased aortic wall stiffness) by 6 months of age. This time point is equivalent to 30-35 human years and was compared to middle-aged 12-month-old CTRL mice, to evaluate the MFS-associated accelerated vascular aging phenotype. These MFS mice are bred on a *C57BL/6* background, making *C57BL/6* mice the appropriate controls for this study. Male and female CTRL and MFS mice were utilized to investigate sex as a biological variable. After genotyping via tail snip and PCR testing, a total of 60 mice were randomly assigned to naïve 6M-old CTRL and MFS (males & females), and 12M-old CTRL mice (N=10/group/sex) and utilized for all measures described.

### High-resolution high-frequency ultrasound imaging

*Vevo®2100* ultrasound system (VisualSonics, FUJIFILM, Toronto, Canada) was utilized to conduct evaluation of functional and structural properties of the aorta, pulmonary artery, coronary artery, and posterior cerebral artery (PCA) along with aortic valves and cardiac function and structure in the same experimental mice for all measures. Experimental Mice (N=10/group/sex) were subjected to *in vivo* ultrasound imaging equipped with a transducer with central frequency of 40 MHz, focal length of 7.0 mm, and a frame rate of 557 fps, at 6M or 12M of age.

The sample size for experimental groups was calculated based on desired endpoints where power was set at 90% and α=0.05. The Mean ± SD were utilized from our and other previously published data to reach a 20% reduction in aortic root dilation (as evident by using ultrasound imaging) in MFS mice in response to intervention (N=10/group/sex).^12,13,62^

Mice were anesthetized with 3% isoflurane and 2L/min 100% oxygen in an induction chamber and kept anesthetized using a nose cone at 2% isoflurane and 1L/min 100% oxygen, while on a heated and maneuverable platform and body temperature monitored via rectal probe; anesthetized state was confirmed by loss of righting reflex and toe pinch. Throughout the procedure, heart and respiratory rate were monitored via an electrocardiogram (ECG) by securing the limbs to ECG electrodes embedded in the heated platform. Hair was removed using a depilatory cream at the sight of imaging (top of the head and chest). All ultrasound imaging data were analyzed using the *Vevo LAB* software by two investigators blinded to age, sex, and genotype during analysis, and the data was reported as average of the two investigators.

### Aortic root diameters and aortic pulse wave velocity measurements

Using a MS550 transducer and B-mode aortic arch view, we measured aortic root diameters at three different regions of the aortic root (aortic annulus, sinuses of Valsalva, and sinotubular junction) in 6-month-old male and female CTRL and MFS mice, and sex-matched 12-month-old CTRL mice. The aortic annulus was measured as the diameter of the transition point between the left ventricle and aortic root. The sinus of Valsalva was measured parallel to the aortic annulus at the largest width of the aortic root. The sinotubular junction was measured parallel to the other diameters and at the junction between the aortic root and the aortic arch (**Fig. 1A & 1B**).^62,63^

We also measured aortic pulse wave velocity (PWV), as a reliable indicator of aortic wall stiffness. As explained in our previous report, aortic peak velocities from the ascending and descending aorta were measured in PW Doppler aortic arch view.^62,63^ The aortic arch distance (mm) was traced along the central axis of the aortic arch at the time of imaging such that the ascending and descending aortic peak velocity coincide with the aortic arch distance (**Fig. 2A**). This distance was measured in B-mode. Transit time, or the time for the pulse wave to travel the aortic arch distance, was calculated as the average of ten consecutive and replicable descending aortic peak velocities minus the average of ten consecutive and replicable ascending aortic peak velocities (**Fig. 2B & 2C**). The PWV was calculated as 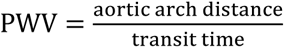.^62,63^

### Coronary and pulmonary peak blood flow velocity measurements

To assess coronary and pulmonary peak blood flow velocity, a MS550 transducer was utilized. These measures were taken in parasternal long axis (PLAX) view in B-mode to visualize the heart. Color Doppler mode was activated to assess blood flow and identify the arteries of interest. The pulse-wave Doppler sample volume cursor was positioned over the coronary artery and then the size and angle were adjusted, and velocity measured. The same was done to assess the pulmonary artery outflow. The velocity waveforms were analyzed as the average of five consecutive and replicable peaks per artery per animal by two independent investigators. All waveform measurements were represented as positive values.

### Posterior cerebral artery peak blood flow velocity measurements

To assess PCA peak blood flow velocity, the MS250 transducer was employed. This measure was completed first such that the mouse was anesthetized for no longer than 5-10 minutes to avoid hyperoxic effects on cerebral blood flow. In B-mode, a coronal section of the mouse cranium was visualized, and color Doppler mode activated to visualize posterior cerebral artery blood flow (**Fig. 5A**). Utilizing pulsed-wave Doppler, the size and angle of flow was adjusted to meet the target artery and a waveform generated (**Fig. 5B**). Five consecutive and replicable peak measurements were analyzed and averaged by two blinded investigators per animal, where the reported values are the average of the investigators.

### Cardiac parameter measurements

Cardiac parameters determined from the left ventricular trace were obtained as previously described.^62,63^ Using the MS550 transducer, the parasternal short axis view of the left ventricle (LV) was visualized in B-mode. The M-mode cursor was then positioned in the middle of the LV and a cross-sectional view of the LV was generated (**Fig. 6A**). During analysis, the LV trace function was utilized to evaluate the functional parameters of heart rate, stroke volume, cardiac output, ejection fraction, fractional shortening, and LV mass (**Fig. 6B**). Three contraction cycles (systole & diastole) were measured per animal and analyzed by two independent blinded investigators and the averaged results are reported.

### Mitral valve measurements

Mitral flow velocity was measured using the MS550 transducer in apical four chamber view. Using color doppler, mitral flow was visualized, and PW doppler cursor size and angle adjusted to generate the velocity waveform. Mitral flow velocity including both early (MV E) and atrial (MV A) velocities were acquired (**Fig. 7A**). During analysis three consecutive and reproducible inflow velocities were assessed for the MV E and MV A velocities for each animal as analyzed by two blinded independent investigators. The E/A ratio of these velocities was also reported to evaluate mitral valve function.^63^

### Statistical analysis

All graphs, curves, and each statistical significance were determined and created with GraphPad Prism software. Two-way ANOVA test was performed on data sets with two independent variables (e.g., genotype & sex). One-way analysis of variance (ANOVA) test was performed on data sets with one independent variable (e.g., phenotype). Due to the exclusion of 12M-MFS cohorts, Two-way ANOVA tests could not be performed on data sets of 6M-CTRL, 6M-MFS, and 12M-CTRL. Our assessment evaluated the phenotypic similarity and differences between these cohorts thus 6M-MFS and 12M-CTRL mice were evaluated for the independent variable of phenotype. Tukey post-hoc was performed on ANOVA tests. All data sets were tested for outliers utilizing ROUT (Q=1%) and if determined were excluded and evaluated for Gaussian distribution to test for normality of residuals. Significance was determined as a *P*-value ≤ 0.05. *P*-values are reported in the graphs and raw data can be accessed by contacting the corresponding author. Personnel were blinded to genotype, age, and sex during analysis. Phenotypic differences induce difficulty with blinding during ultrasound imaging, thus all images were de-identified, re-coded, and blinded for analysis by two independent investigators. Pre-determined welfare exclusion criteria include removing any mouse from the study with visible wounds requiring veterinarian intervention or vocalization of pain which cannot be managed. No animals required exclusion due to welfare concerns. In some circumstances, not all measurements could be assessed for each animal due to difficult imaging such as “sternum shadow” interferences, leading to inaccurate measures that were excluded from analysis.

## Author Contributions

TMC designed experiments, collected, and analyzed data, composed data figures, and wrote the manuscript. MEB helped with data collection and independent analysis. ME and TCT supervised and funded the project, helped with experimental designs, reviewed the data, revised and edited the written manuscript. All authors approved the final version of the manuscript.

## Data Availability Statement

The data sets generated for this study are available upon request to the corresponding author.

## Ethics Statement

Animal care was conducted according to the National Research Council Guide for the Care and Use of Laboratory Animals and the Guidelines for Animal Experiments and the Institutional Animal Care and Use Committee. All experiments were in compliance with ARRIVE guidelines. The animal study was reviewed and approved by the Institutional Animal Care and Use Committee [IACUC protocols AZ-3006, AZ-2989, & AZ-2936] at Midwestern University-Glendale, Arizona.

